## Supplementary Material for "Bipartite network analysis of ant-task associations reveals task groups and absence of colonial daily activity"

**Table S1.** *Diacamma* sp. colony information. The number of broods (egg pile, larvae and pupae) and colony size indicate the number of brood and ants in the colony at the start of the recording.

| Colony ID | Num. of ants | Brood | Num. of egg pile | Num. of larvae | Num. of pupae | Availability of data on Queen | Colony size | Num. of lost ants |
|---|---|---|---|---|---|---|---|---|
| a | 33 | egg / larva / pupa | 10 | 5 | 10 | - | 37 | 4 |
| b | 41 | pupa | 0 | 0 | 13 | + | 43 | 1 |
| c | 78 | absent | 0 | 0 | 0 | - | 88 | 10 |
| d | 104 | egg / larva / pupa | 0 | 18 | 3 | + | 112 | 8 |
| e | 121 | larvae | 0 | 20 | 0 | - | 165 | 44 |

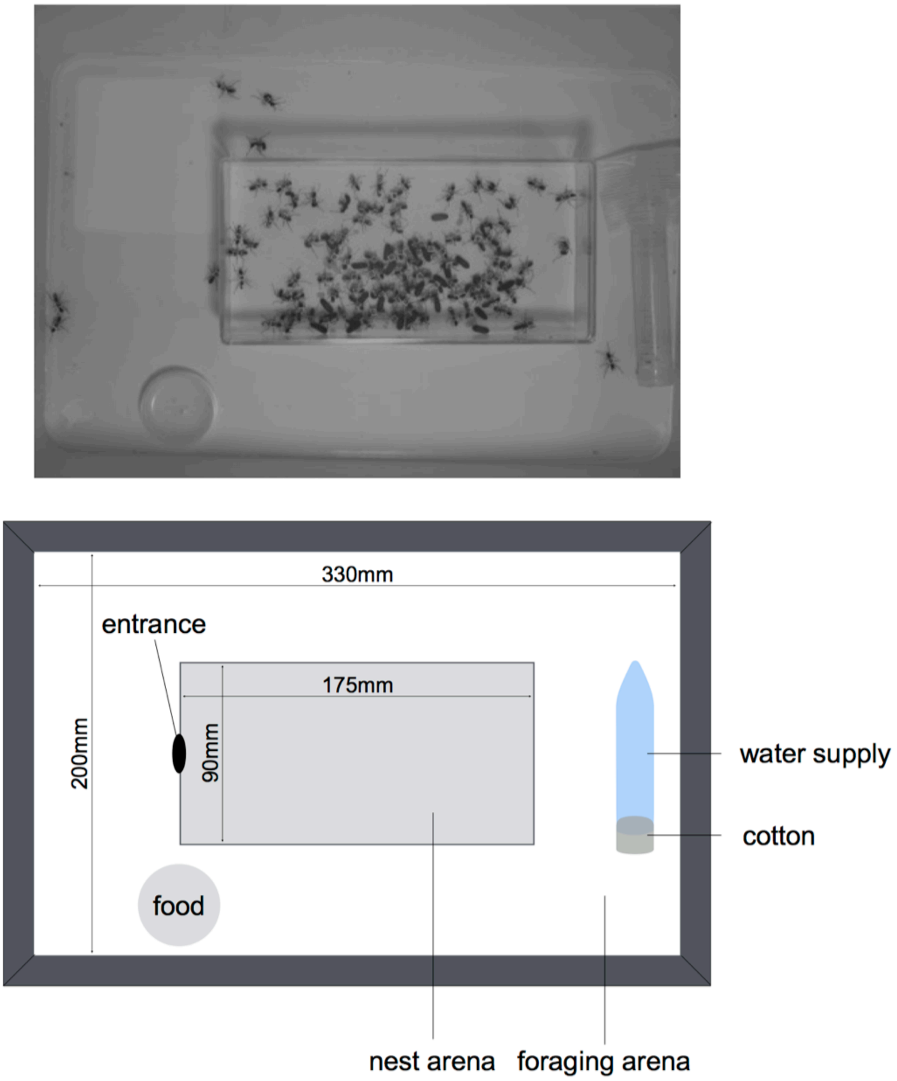


**Figure S1.** Experimental setup. The artificial nest was a square plastic box that was filled with a moistened plaster.

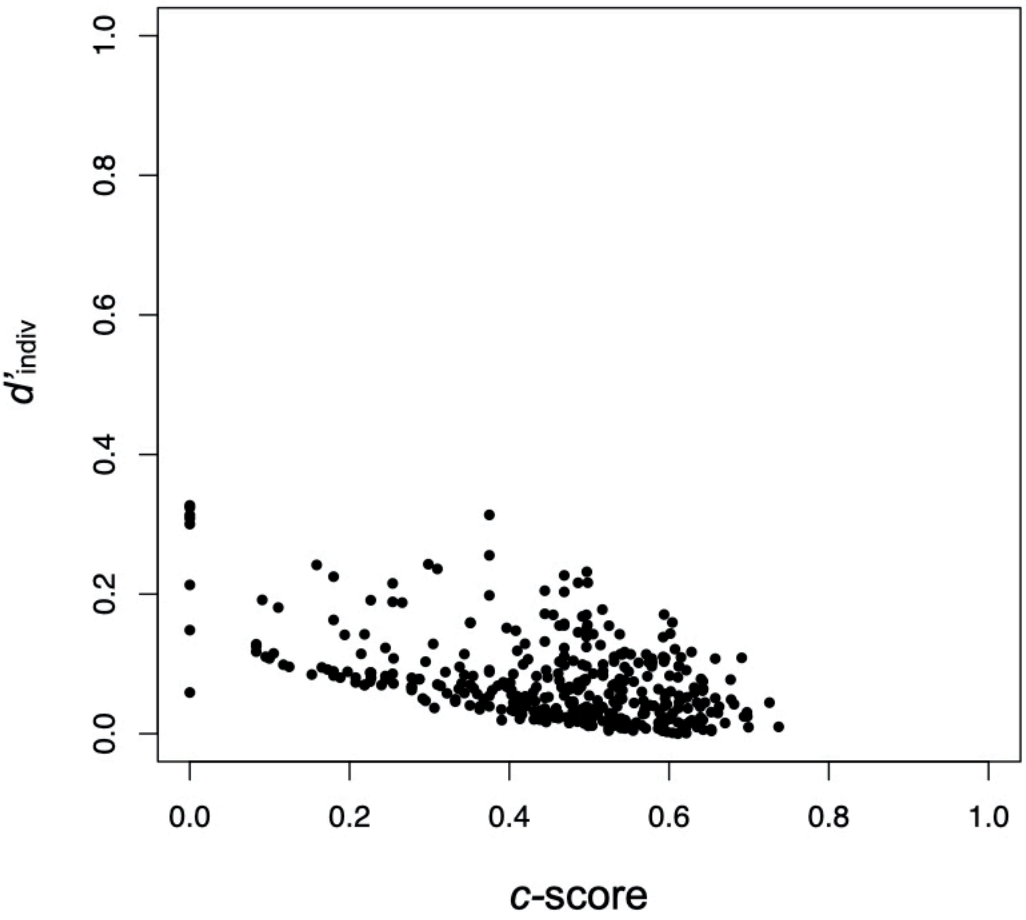


**Figure S2.** Relationship between $d'_{\text{indiv}}$ and *c*-score. There is a significant correlation between two metrics (correlation coefficient -0.48).

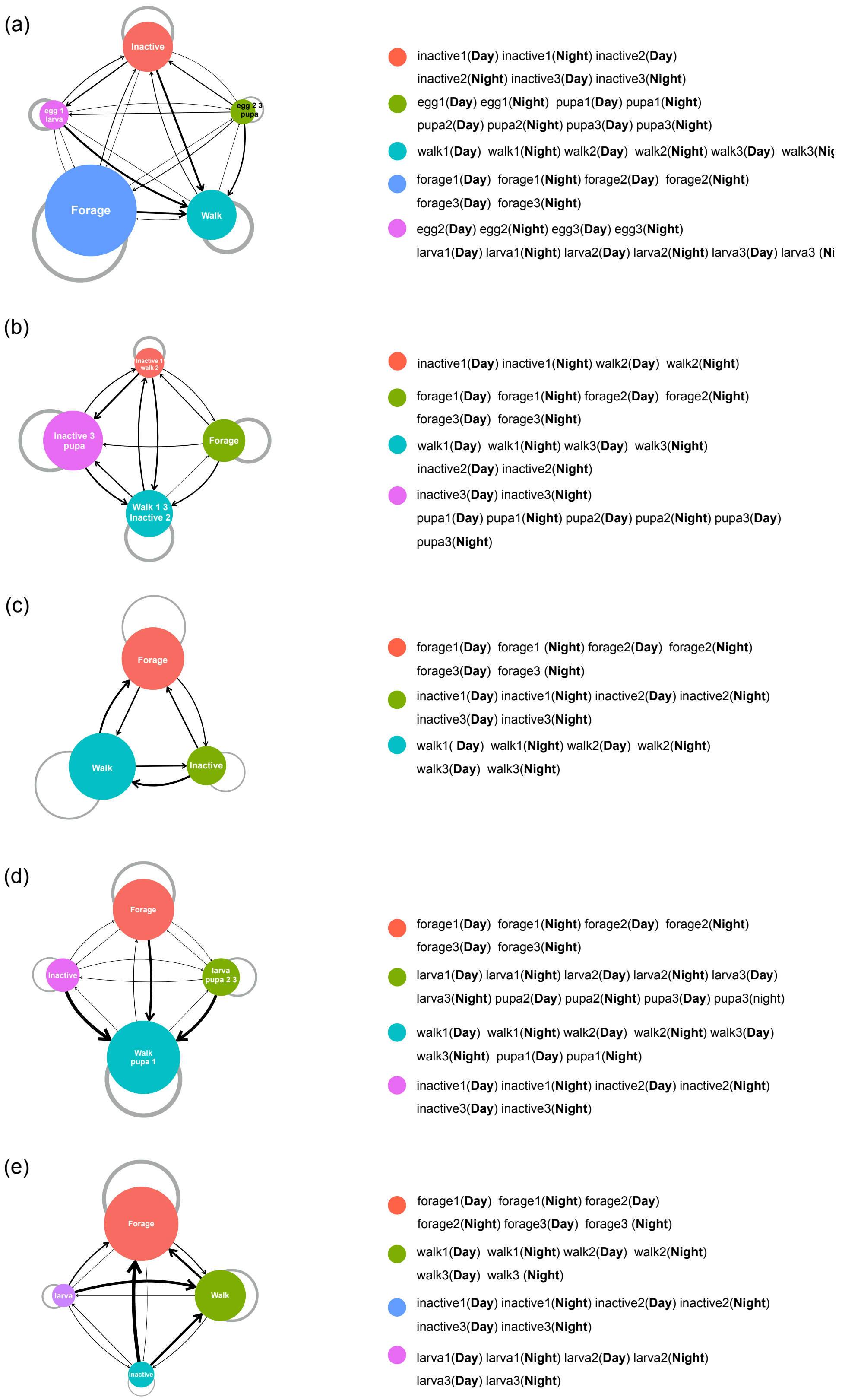


**Figure S3.** Networks of modules in 5 *Diacamma* sp. colonies (a to e) when separated data into Day and Night. The node represents modules containing individuals and behaviour. The labels on the nodes indicate the behaviour contained in the module. The size of the nodes is weighted by the number of individuals belonging to the module. The links have a direction and width as the mean weight from ant individuals belonging to module *i* to behaviour belonging to module j ($q_{ij}$). The width of grey self-links indicates the mean weight from ant individuals belonging to module *i* to behaviour belonging to the same module *i* ($q_{ii}$). Right panels indicate behaviour with day information (day 1, day 2, or day 3) that belong to each module.

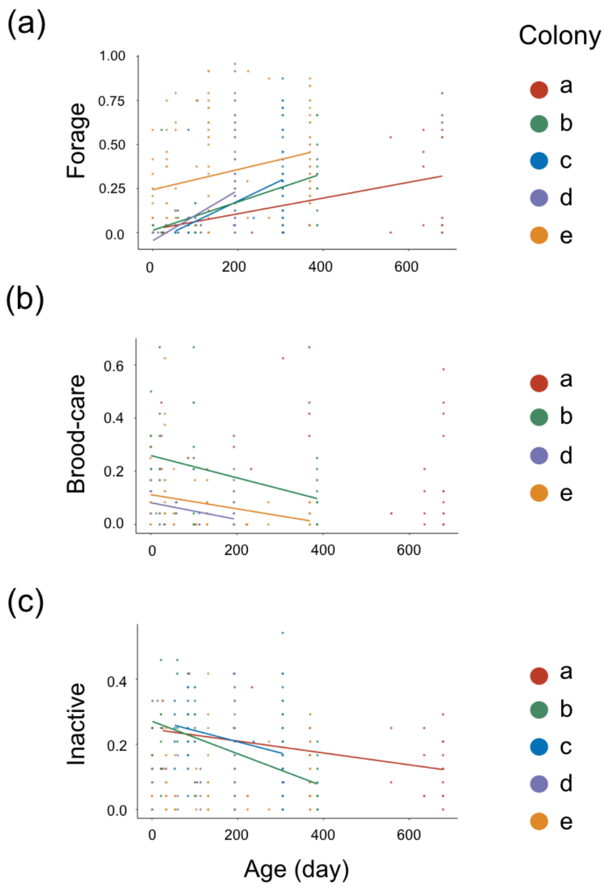


**Figure S4.** The proportion of *Diacamma* sp. behaviour (a) forage, (b) brood-care, and (c) inactive with respect to the age of an individual. Each point indicates an individual and the colour represents the colony. No data about broods is in colony c because the colony c did not have broods.
